# Electrophysiologically Targeted Biopsies Reveal the Transcriptional Landscape of Focal Epilepsy

**DOI:** 10.64898/2026.03.06.708552

**Authors:** Ashwin Viswanathan, Molly Murch, Abby Brand, Julia L Furnari, Nathaniel W Rolfe, Archana Yadav, Clara H Stucke, Aayushi Mahajan, Juncheng Li, August Kahle, Misha Amini, Tristan T Sands, Osama Al-Dalahmah, Jeffrey N Bruce, Brian J A Gill, Neil A Feldstein, Brett E Youngerman, Guy M McKhann, Vilas Menon, Peter Canoll, Melodie Winawer, Catherine A Schevon

## Abstract

Up to 30% of patients with epilepsy have intractable seizures, yet the mechanisms of focal ictogenesis remain unclear. Tissue involvement in ictal regions is heterogeneous, with different regions playing distinct roles in ictogenesis, seizure propagation, and resistance to spread. These roles are reflected in electrophysiologic differences between the seizure focus and the ictal penumbra, where evidence of synaptic spread is present but excitatory firing is constrained by largely intact inhibition. Investigation of the disruption of the normal interplay between excitatory and inhibitory activity, thought to underlie ictogenesis across a range of epilepsy etiologies, is limited by network complexity and cellular heterogeneity in human tissue samples. In this study, we relate cellular and molecular alterations to excitatory-inhibitory disruption in network dynamics defined by electrophysiologic features. This work may aid in the identification of clinically relevant tissue biomarkers and support novel preclinical therapeutic approaches for treatment-resistant focal epilepsy disorders.

We developed a novel intracranial EEG guided, MRI-localized approach to sample paired biopsies from 11 patients with drug-resistant focal epilepsy with diverse etiologies, which were then studied using single-nucleus RNA sequencing (snRNAseq) and immunohistochemistry (IHC). EEG recorded from stereotactically implanted depth arrays (sEEG) was used to identify regions of epileptic involvement, based on findings from prior simultaneous clinical and microelectrode recordings. This approach addresses the intrinsic heterogeneity due to etiology and cortical architecture through paired, within-patient comparisons.

We identified distinct cell-type specific transcriptional signatures that differentiate cellular populations in the seizure focus and ictal penumbra in intractable focal epilepsies. Our findings provide a link between tissue composition and gene expression that correlate with electrographic features in a heterogeneous seizure landscape. Our findings support common pathways of seizure generation and spread that are conserved across disease etiologies. Relative depletion of interneuron populations in the seizure focus supports the hypothesis of disrupted inhibition as a driver of epileptiform activity in the seizure focus. The enrichment of plasticity-associated gene signatures in the penumbra suggests a complex interaction of these regions with the seizure focus, as well as the role of the penumbra in enabling or limiting seizure expansion. This study provides a novel methodology for tissue sampling in epilepsy and uncovers biologically relevant tissue signatures that provide grounds for future work in targeting cellular and molecular alterations present in focal epilepsies.

## Introduction

While surgical intervention has shown efficacy in treating neocortical focal epilepsies, a significant number of patients who undergo epilepsy surgery continue to have medically intractable seizures, leading to increased morbidity and mortality. Extensive research continues to be focused on the mechanisms of epileptogenesis and seizure generation, and identification of sEEG biomarkers to guide surgical treatment. To date, there have been limited efforts to map the molecular and cellular landscape of epileptic brain regions to uncover the long-term consequences of seizure activity in involved tissues,^1–9^ and to highlight pathological mechanisms that may be exploited to refine therapeutic medical or surgical treatments.

This study tests for differences in cell-type specific mRNA expression between two key brain territories involved in focal neocortical seizures: the seizure focus, the region from which focal seizures are thought to arise, and the ictal penumbra, the surrounding brain regions subjected to strong excitatory synaptic currents during seizures but which lack electrophysiological hallmarks of local seizure invasion or paroxysmal depolarizations.^10–14^ These dynamics reflect the collapse of inhibitory restraint in the seizure focus, whose hallmark electrophysiologic feature is a massive increase in synchronized local excitatory activity that drives seizure activity. Beyond this boundary, inhibitory tone is largely preserved, restraining excitatory neuronal firing despite the presence of pathologically strong excitatory drive.^3,15–19^ This results in a heterogeneous neuronal activity pattern, instead of the synchronous neuronal firing characteristic of the seizure focus. In the penumbra, EEG ictal rhythms can be prominent due to local response to strong excitatory currents. As long as inhibition remains intact, however, the penumbra does not drive seizures and in fact may resist local seizure invasion.^19,20^ Comparisons between these regions have been limited to *ex vivo* electrophysiological or calcium imaging observations during seizures in acute animal models or in human tissue recorded simultaneously with microelectrodes. We seek to identify the cell-type specific differences between the two territories by probing their transcriptional signatures. Our goal is to uncover conserved, biologically relevant trends that distinguish the seizure focus from surrounding tissue across focal epilepsy etiologies, including developmental and acquired conditions.

Disturbance of the interplay between excitatory and inhibitory neuronal subtypes has been implicated in ictogenesis.^21–23^ Physiological cortical architecture involves balanced activity of excitatory neurons and multiple distinct inhibitory interneuron subtypes. Recent work has identified a key role of interneuron subtypes in physiological cortical function and in constraining epileptiform activity in the setting of epilepsy.^24–27^ Dysfunctional inhibition, which occurs with and contributes to changes in excitatory axonogenesis and synaptic formation in network circuitry, has been hypothesized as a leading mechanism for ictogenesis and seizure spread in acquired epilepsies.^28^ Some hypotheses of driving factors for this inhibitory collapse include chloride dysregulation in pyramidal cells contributing to GABAergic dysfunction,^5,29–34^ abnormal interneuron function,^35–38^ and cell-type specific alterations to interneuron activity.^39^ An alternative hypothesis has recently been proposed that suggests interneuron activation can “jump-start” seizures via an excitatory rebound mechanism, thus forcing synchronized excitatory firing.^40,41^ Preclinical experiments have implicated this collapse of inhibitory drive, some with a documented loss of interneuron function associated with focal epilepsy, to be a key contributor to seizure propagation^12,15,42–45^ With this context, the complexity of involved cell-types necessitates single-cell resolution to identify cellular and molecular tissue features that distinguish these regions of differential excitatory-inhibitory function.

Electrographic localization via invasive EEG has been used both clinically and in prior research studies to identify regions of seizure involvement.^46^ Few studies have directly used this localization to probe the gene signatures of involved cells to assess for changes in tissue state related to focal epileptiform activity. In this study, we thus employ a novel MRI-localized, electrographically-guided methodology to identify the seizure focus and penumbra and directly probe tissue-based signatures of these distinct territories. We provide a targeted approach to acquiring and profiling tissue from distinct MRI-localized, EEG-guided biopsies from patients undergoing epilepsy surgery (**Fig. 1)**. We employ this novel methodology to pair biopsies from the seizure focus and ictal penumbra, allowing for within-patient comparison via snRNAseq and IHC. Our analyses provide valuable insight into tissue states in the seizure focus and reactive effects in the ictal penumbra, and foster development of novel therapeutic strategies based on these molecular and cellular findings.

**Figure 1.**
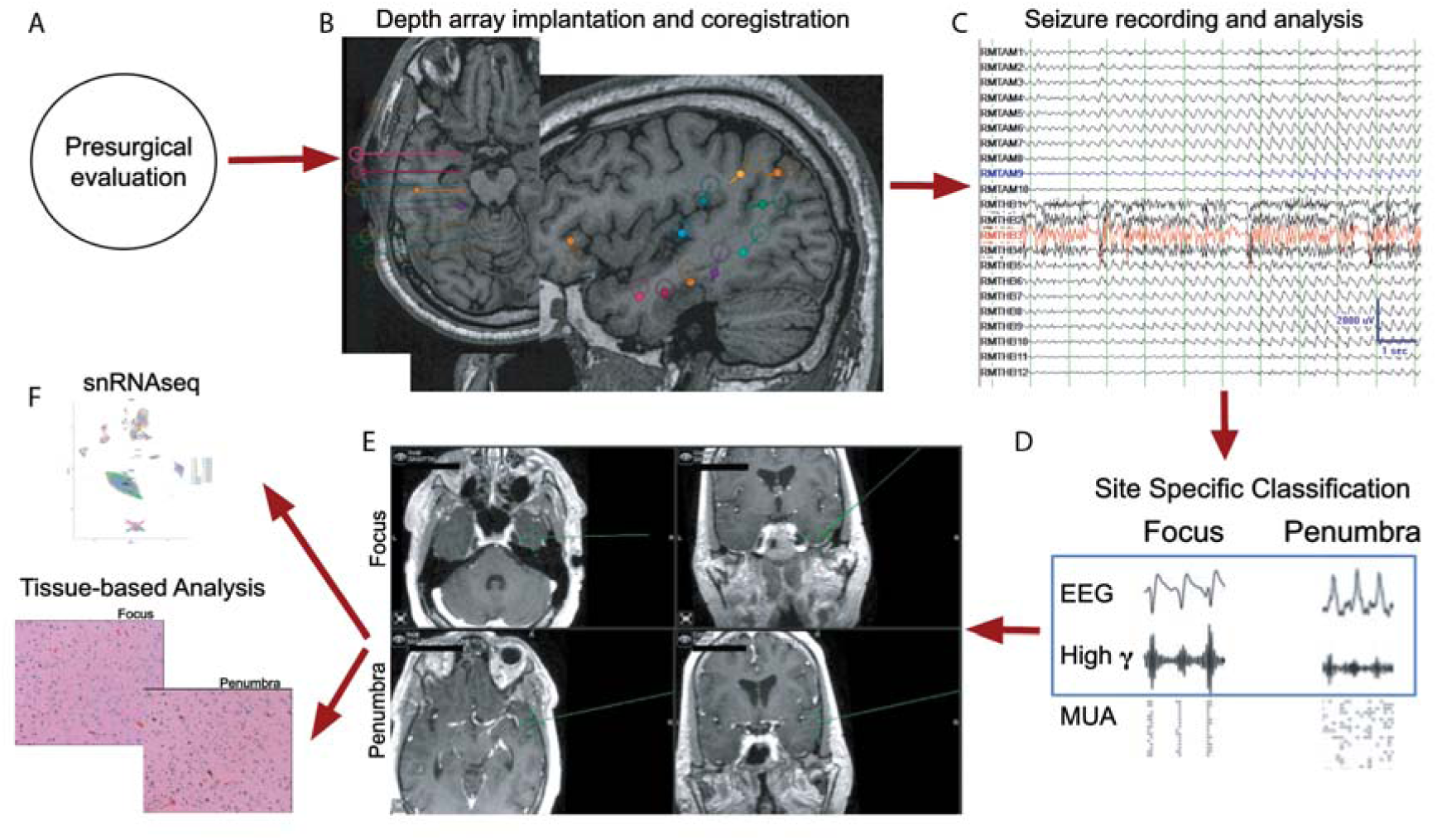
Sampling and Labelling Workflow. Exemplary workflow using imaging and data for a single patient. **(A)** Patients undergo presurgical evaluation for medically intractable focal epilepsy and are recommended for invasive sEEG monitoring to delineate the epileptic brain region. **(B)** After depth array implantation, electrodes are localized to the pre-operative volumetric MRI, and long term recording is conducted to capture spontaneous, typical seizures. **(C)** For study purposes, the EEG and high gamma components are visually assessed to identify the seizure focus (red) and penumbra (blue) territories. **(D)** Sample recordings from previously published work from our group include multiunit activity raster plots demonstrating the sharp contrast in neural firing patterns underlying the two regions.^14^ Our dataset includes only EEG and high-gamma components (blue box). MUA activity is shown to demonstrate underlying activity representative of the two territories. **(E)** Biopsies are taken from both territories, with sites documented using a probe co-registered to the volumetric MRI. **(F)** Samples are then processed for snRNAseq and IHC.

## Materials and methods

### Patient Enrollment

#### Enrollment and Inclusion Criteria

An overview of the patient recruitment process and criteria is shown in **Fig. 2**. This study screened 49 adult and pediatric patients with medically intractable focal epilepsy syndromes who underwent invasive EEG-guided neocortical brain resection as a therapeutic procedure to control seizures at Columbia University Irving Medical Center between 2014-2025. Of this larger cohort of patients, 24 had brain samples collected as part of this study. Clinical determination of eligibility for focal resection was determined by CUMC’s standard clinical practice, incorporating seizure semiology, electroencephalographic (EEG) data, structural MRI imaging, functional imaging (positron emission tomography, single photon emission computed tomography), and clinical consensus at epilepsy surgery conferences. Patients who had a focal onset epilepsy diagnosis and surgical implantation of electrodes with a plan for subsequent, potentially curative resection were considered as potentially eligible.

**Figure 2.**
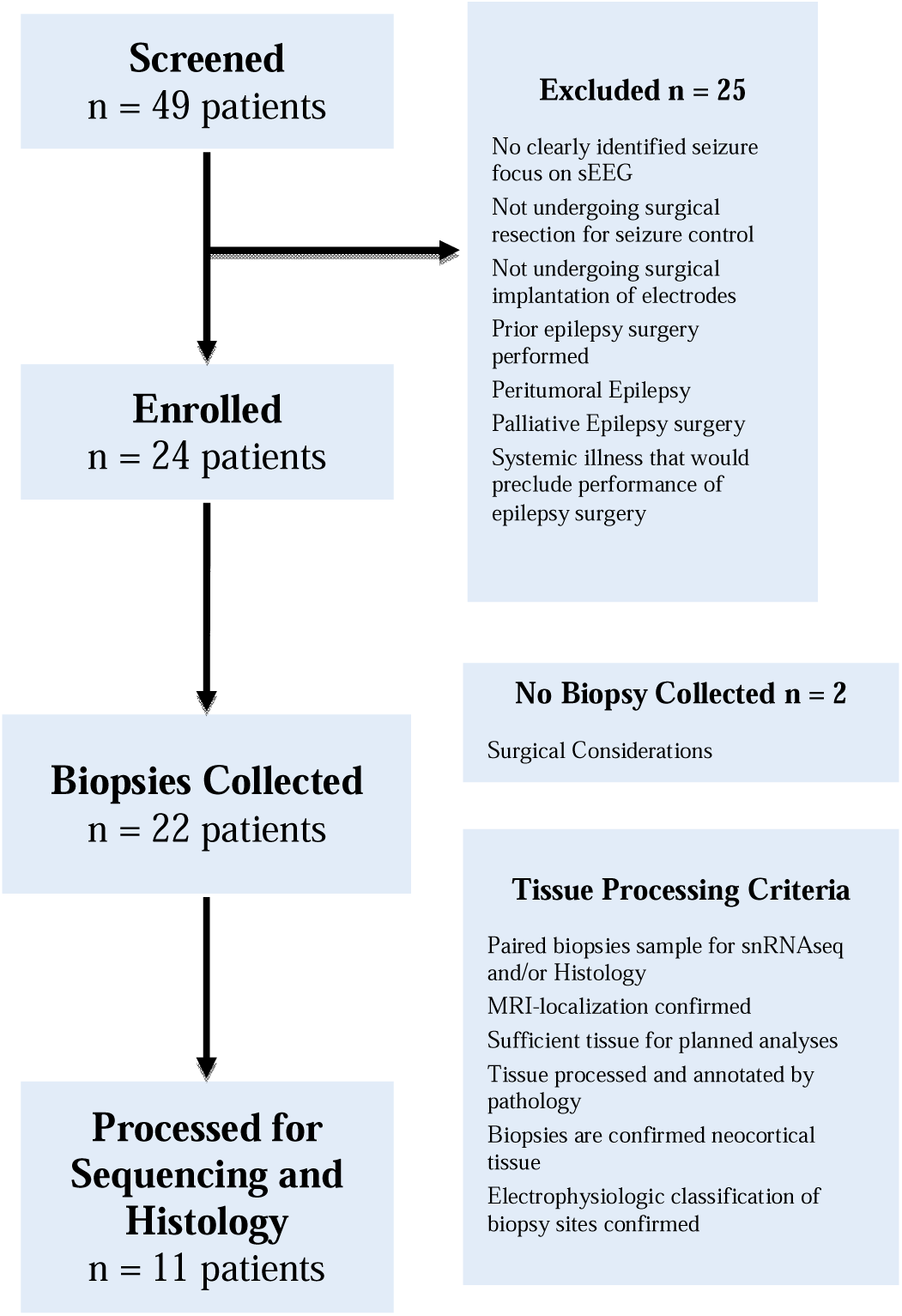
Patient Recruitment Flow Chart. Flow chart indicating number of patients included at each stage of the study pipeline and describing criteria for exclusion/inclusion. 49 patients were initially screened, of which 24 were enrolled. 11 patients were ultimately included for snRNAseq and histologic analysis.

Prior to surgical resection, patients additionally underwent invasive EEG monitoring with stereographically implanted depth electrodes (sEEG), with array trajectories and locations determined according to clinical needs.^47^ Following invasive monitoring, all cases were discussed at our center’s multidisciplinary surgical case conference to reach consensus on a therapeutic plan, as per standard clinical practice. Patients who were recommended for therapeutic resection of epileptic tissue, determined based on the preceding presurgical and invasive diagnostic studies, were then considered eligible for study screening.

#### Patient Screening and Consent

From this population, patients were excluded if they declined or were unable to undergo brain resection due to systemic illness that would preclude favorable outcomes from surgical intervention, had prior epilepsy surgery including palliative epilepsy surgery (e.g. Rasmussen’s), had peritumoral epilepsy resulting from metastatic or primary brain neoplasm (except for WHO grade 1 tumors), or whose seizure focus could not be localized within a circumscribed structural brain area or ∼2 cm brain region, which is less than the average extent of seizing brain in neocortical focal epilepsy cases.^48^ Patients undergoing laser interstitial thermal ablation were excluded due to lack of tissue availability. Due to likely differences in cell-type specific expression profiles in mesial temporal structures such as hippocampus in comparison to neocortex, patients for whom seizures were found to originate solely in hippocampus or amygdala were also excluded.

Informed consent was obtained for all patients under an approved Columbia University Irving Medical Center Institutional Review Board protocol. Patients of all ages were eligible, assuming that they had been deemed eligible for surgical treatment of epilepsy. For minor subjects, parental informed consent was obtained, along with assent for children aged 14–17. All 24 enrolled patients provided verbal consent via phone and written consent following a review of the study’s consent form, which outlined confidentiality practices (data deidentification, assignment of a unique identification code, and password protected databases) and data sharing expectations (required sharing of genetic data in anonymized form to an NIH-managed database and possibility of sharing of deidentified data with other researchers for epilepsy research purposes).

### Pre-Operative Planning

#### sEEG Implantation and Pre-Operative Monitoring

Arrays of 8-16 contacts each were utilized (PMT Corp., Chanhassan, MN or Dixi Medical USA Corp., Oxford, MI), with electrode positioning with respect to brain structures following planned trajectories using ROSA (Zimmer Biomet, Warsaw, IN) and verified by co-registration of the post-implant CT with pre-operative volumetric MRI at the time of implant (Brainlab, Westchester, IL). Long-term video EEG monitoring was then conducted using a Natus Quantum system with 2 kHz sampling rate and 0.1-512 Hz hardware bandpass (Natus Medical Inc., Middleton, WI) to capture interictal discharge populations and typical seizures. Following passive recording, most patients then underwent electrical stimulation mapping procedures (conducted by CAS) to identify sites with critical language, motor, and/or vision function, and sites from which seizures could be triggered correlating with the patient’s typical semiologies. This is a routine procedure at our center that has been shown to predict post-resection seizure control.^49,50^

#### Localization of Seizure Territories

The seizure focus was localized based on passive sEEG seizure recordings, with seizure onset sites first determined by the clinical team with later confirmation by CAS and the treating neurosurgeon (GM, BY, or NF) (**Fig. 1**). Seizure onset was defined as the sites in which the earliest departure from usual interictal patterns was seen, represented by onset of low-voltage fast activity, with evolution in frequency, amplitude, and morphology to a sustained, full seizure. As the presence of ictal waveforms does not suffice to distinguish the seizure focus from the surrounding penumbra, additional supporting information was used to make a final determination of the location of the seizure focus.^51^ This included diagnostic studies from the presurgical evaluation phase, including imaging findings and seizure semiology, characteristic seizures provoked by electrical stimulation, and the presence of high gamma activity modulated by the ictal rhythm, a method that has been validated using surgical outcome studies^48,52^ and evidence from time-locked microelectrode recordings demonstrating single and multi-unit firing patterns during the initial phases of seizures.^53,54^

### Patient Biopsies and Tissue Processing

#### Sampling and Tissue Handling

Patients undergoing therapeutic brain resection who received diagnostic invasive sEEG were selected for the study. Brainlab neuro-navigation was utilized intraoperatively to localize brain sites identified by sEEG, and to provide a visual record of biopsy sites for post-hoc confirmation (**Fig. 1**). Prior to carrying out therapeutic resection, surgeons took biopsies from selected sites, with sites chosen via pre-procedure review. Brain samples were received by neuropathology and released for research use after initial evaluation.

Biopsies were full-thickness and included all cortical layers. Radiographic localization was confirmed using intraoperative stereotaxic instrumentation paired with guided MRI sequences. If enough tissue was recovered, samples from each region were bisected in the operating suite along an axis perpendicular to the pial surface. One portion was processed for snRNAseq and was flash frozen in liquid nitrogen. A second portion was processed for histological and IHC analyses and was placed in 10% formalin. We obtained paired samples from 11 patients that were included for snRNAseq, and paired samples from 9 patients for IHC analysis.

#### Tissue Immunohistochemistry

Tissue biopsies intended for histological analysis were then embedded in paraffin. 5-micron tissue sections were then stained with hematoxylin and eosin. IHC staining on tissue blanks was performed using rabbit anti-NeuN antibody (Cell Signaling; 1:500), rabbit anti-GFAP (Dako; 1:5000), rabbit anti-Iba1 (Cell Signaling; 1:200), mouse anti-Parvalbumin (Sigma; 1:750), and rabbit anti-RORB (1:500) antibodies and counterstained with hematoxylin. Each slide was scanned using a Leica SCN 400 digital slide scanner at 40x magnification. Images were analyzed in QuPath image analysis software. Regions of each tissue were scored by a blinded neuropathologist to specify regions of cortical tissue. A labeling index was then calculated for each stain by counting cell positivity as a fraction of total cells labelled via hematoxylin. For neuronal staining (NeuN, PV, RORB), we limited analysis to only include cortical regions as identified by a trained neuropathologist.

### Single-Nucleus RNA Sequencing and Quality Control

snRNAseq was performed similarly to previously published methodology.^55^ Nuclei were isolated from flash-frozen biopsy samples in accordance with established protocols described previously.^55–57^ Libraries were prepared using Chromium Next GEM Single Cell 3’ Reagent Kit v3.1 (PN 120237), with Chromium Single Cell A Chip Kit, 48 runs (PN 120236). Target cell recovery was at least 10,000 cells per sample for both focus and penumbra samples. The final number of extracted nuclei was calculated using DAPI as a marker and calculating an average of three counts on a Countess (Thermo Fisher). The index plate used was 10X Dual Index Kit TT Set A (PN 1000215). Sequencing was done using 10X Chromium v2 chemistry, and reads were aligned to the GRCh38 human genome with Refseq v93 transcriptome annotation using Cellranger v6.0.1 for quantification of transcripts belonging to each nucleus. The quantified count matrices for each sample were then run through CellBender v3 using –expected-cells=5000, --total-droplets-included=25000, and all other parameters set at default values.

Output matrices from CellBender were then imported into R v4.0.1 and analyzed using the Seurat v4 package for clustering and cell type identification. After concatenating all CellBender counts matrices, only nuclei with >=500 total non-mitochondrial, non-pseudogene, non-antisense gene and non-gene-model Unique Molecular Identifiers (UMIs), and <=5% mitochondrial UMIs were retained. The remaining nuclei were normalized and scaled (using the LogNormalize and ScaleData functions with default parameters), and variable genes were identified using the FindVariableFeatures function with default parameters. Individual samples were then batch-corrected and integrated using the Harmony workflow with PCs 1-30, and Harmony dimensions 1-30 were used to create a k-Nearest Neighbors graph (FindNeighbors function with k.param=8) and a UMAP for visualization. Finally, clusters were identified from the k-Nearest Neighbors graph using the default Louvain algorithm in the FindClusters function with resolution = 1.

From this initial analysis, all clusters were assigned to broad cell classes by examining the expression of the following genes: SNAP25, RBFOX3, SLC17A7, GAD2, SLC32A1, OLIG2, MOG, MAG, PDGFRA, AQP4, GFAP, FGFR3, ALDH1L1, CD24, AIF1, TMEM119, C1QA, CTSS, MRC1, CLDN5, RGS5, PDGFRB, CD3E, CD8A. Clusters were manually assigned to the following classes: Glutamatergic Neurons, GABAergic Neurons, Oligodendrocytes, Oligodendrocyte Precursor Cells, Astrocytes, Myeloid Cells, T-Cells, and Vascular Cells. Clusters with markers for more than one broad class were assigned as doublets and removed from further downstream analysis. Clusters with 90% of cells coming from a single sample were also removed from downstream analysis. This entire process (starting from log normalization and scaling, and subsequent steps using the same parameters) was then repeated for each broad class of cells, with clusters having mixed signatures being removed in 2 separate iterations. After the second iteration of mixed signature cluster removal, individual clusters for each broad cell class were remerged across all broad cell classes to generate a new UMAP for visualization; note that cluster identities from the subclustering were retained, and the entire data set was not reclustered as a whole. This process then resulted in a broad class and subcluster annotation for every nucleus that was retained.

For further annotation of subclusters, especially neuronal subclusters, the entire data object was saved as an h5 AnnData object and passed into the Allen Institute’s MapMyCells online server; the annotations output from the server were then used to label neuronal subclusters by their Allen Institute reference atlas supertype, with additional numerical distinctions where needed. Glial subclusters were labeled numerically.

For differential cluster proportion abundance between focus and penumbra samples, a nonparametric paired-sample test (Wilcoxon signed rank test) was performed for each cluster separately to determine significance (p < 0.1), using its proportion of the total broad class as the paired variable for each donor. For one donor with multiple penumbra samples, the mean proportion value was used in the test.

### Consensus Single-Cell Hierarchical Poisson Factorization

To identify co-expression patterns that vary between cells in the focus and penumbra, we generated a consensus scHPF model^58^ for merged focus and penumbra samples within each cell type (excitatory neurons, inhibitory neurons, microglia) across donors. This process was performed using previously described methodology^59^. Batches were down-sampled so that the median unique molecular identifier (UMI) number per cell in different batches was equivalent to the lowest median UMI count among our samples. Testing and training subsets were created for each cell type. For excitatory neurons, 500 cells were taken from each focus and penumbra group across donors for the testing dataset, and for inhibitory neuron and microglial subsets, 200 cells were used. Of note, for quality control in our microglial analysis, one patient was excluded due to an imbalance between focus and penumbra samples in cell identification counts for the testing dataset. Following testing allocation, the remaining cells for each cell type subset were used to create training data objects. A training object was generated using the scHPF prep command with a whitelist (−w) of genes detected in at least 1% of cells (min-cells, m=0.01) across the final downsampled training matrix. Testing objects were generated using the scHPF prep-like command. Consensus scHPF was performed with the following parameters: k = 13–20, t = 5, n = 3, m = 3. The training data object was scored using the final scHPF consensus model and training object gene list generated by the scHPF prep command. This process creates a factor-by-cell matrix with cell scores for each identified factor, a gene list for each factor with contributing genes ranked per factor, and gene scores for each gene in this ranked gene list.

To identify factors that represent biological signals that differ between cells in the focus and penumbra, cell scores for each factor were averaged for each patient sample. Paired analyses were performed to identify factors that exhibited statistically significant differences between focus and penumbra samples via two-tailed paired t-test. For factors that were differentially expressed (p < 0.05), a subsequent Gene Set Enrichment Analysis (GSEA) was performed using established Gene Ontologies of Biological Processes (GOBP) to inform factor contributors. Manual examination of each of these factors and GOBP pathway analysis provided statistically-relevant identities for each factor. For gene ontology analysis, we conducted analysis with biological process annotation, and the Benjamini-Hochberg correction was used to correct p-values for multiple testing. Corrected p-values below a threshold of 0.05 were chosen as significant for annotation results.

### Statistical Analysis

Unless stated otherwise, all analyses were performed using custom scripts in MATLAB, Python, or RStudio. The two-tailed paired t-test was used to test significance unless otherwise noted. For IHC labeling indices and snRNAseq cell counts, the level for statistical significance (α) was set at 0.1. For comparison of factors that were differentially expressed, α was set at 0.05. Mean labeling index ± SEM was used in descriptive statistics.

### Data availability

The data sets are available from the corresponding author on reasonable request.

## Results

### Patient Sample Characteristics

Forty-nine patients met criteria for study screening and 24 patients undergoing epilepsy surgery for intractable focal epilepsy of varying pathologic origin were enrolled (**Fig 2**). 13 of the 24 enrolled patients were male (54%), and patients ranged in age from 2 – 68 years with an average age at enrollment of 30 years. 11 patients whose biopsies were sampled were adults (> 18 years at enrollment). Neuropathologic diagnoses in this population included Focal Cortical Dysplasia (FCD) Type 1a, FCD Type 2a, FCD Type 2b, and generalized gliosis identified on histopathology, including astrogliosis and Chaslin’s gliosis. Of the 24 patients enrolled, biopsies from 22 patients were collected during surgery. Of these, 11 patients were selected for tissue processing according to these criteria: paired biopsies for snRNAseq and/or histologic analyses, confirmed MRI-localization, sufficient tissue for planned analyses, tissue that was processed and annotated through Pathology as confirmed neocortical tissue, and confirmed electrophysiologic classification for tissue samples (**Table 1**).

**Table 1.**

| Subject ID | Age at Seizure Onset (y) | Age at Surgery (y) | Duration of Epilepsy (y) | Site of Surgery | Time from Implant to Resection (d) | Seizure Type(s) | Neuropathologic Findings | Engel Outcome | History of Status Epilepticus |
| --- | --- | --- | --- | --- | --- | --- | --- | --- | --- |
| 1 | 13F | 24 | 11 | F | 24 | FIC, FBTC | FCD 1a | ID | N |
| 2 | 7M | 19 | 12 | O | 80 | FBTC, FIC | Gliosis | IIA | N |
| 3 | 15F | 43 | 28 | T | 265 | FIC, FPC | No diagnostic abnormalities | IV | N |
| 4 | 17M | 21 | 4 | T | 321 | FBTC | Gliosis | IA | N |
| 5 | 15M | 23 | 8 | F | 8 | FBTC | Gliosis | ID | N |
| 6 | 53M | 59 | 6 | T | 124 | FIC, FBTC | Gliosis | IA | Y |
| 7 | 15M | 18 | 3 | F | 18 | FPC | Astrogliosis | IIA | Y |
| 8 | 40M | 41 | 1 | F | 90 | FIC, FPC, FBTC | FCD 2a | IA | N |
| 9 | 17M | 20 | 3 | F | 35 | FIC, FBTC | Gliosis | IA | N |
| 10 | 7M | 48 | 41 | F | 76 | FIC, FPC | Gliosis, FCD 2b | III | N |
| 11 | 38F | 41 | 3 | T | 439 | FBTC | Gliosis | IC | N |

FIC: Focal Impaired Consciousness, FBTC: Focal to Bilateral Tonic-Clonic, FPC: Focal Preserved Consciousness, FCD: Focal Cortical Dysplasia
Patient demographics and characteristics for the 11 patients whose samples are included in the analysis in this study. Patients exhibited a range of intractable focal epilepsy etiologies and neuropathologic findings. Mean age at seizure onset from sampled biopsies is 24.3 (range 7 – 53) years, mean age at time of surgery is 35.2 (range 19 – 59) years, and mean duration of epilepsy prior to surgery is 10.9 (range 1 – 28) years. Average time between sEEG implant and surgical resection in this cohort is 123.3 (range 8 – 439) days. Among enrolled patients, the average age at seizure onset was 16 years, and average duration of epilepsy at enrollment was 15 years. 38% of enrolled patients self-identified as White, 20% as Black, 4% as Asian, and 38% as Other, Unreported, or Unknown. 29% of enrolled patients self-identified as Hispanic or Latino.

### snRNAseq Identifies Cellular Compositional Landscape

Analysis of the single-nucleus transcriptomes were assessed via Uniform Manifold Approximation and Projection (UMAP) clustering. From expression patterns, we were able to identify all major cell populations as distinct clustered populations **(Fig. 3A)**. We also confirmed that samples for each patient **(Supplementary Fig. 1A)** and from both focus and penumbra **(Supplementary Fig. 1B)** were represented in each cell cluster. Cell-types were then identified via gene signatures and subdivided into their specific neuronal subtypes **(Fig. 3B)**. For example, excitatory pyramidal cells were separated by layer-specific markers, with L2-CUX2 expressing cells showing distinct clusters from L4-RORB pyramidal cells. Interneurons were classified by cluster expression of principal markers like PV, VIP, SST, and NDNF.

**Figure 3.**
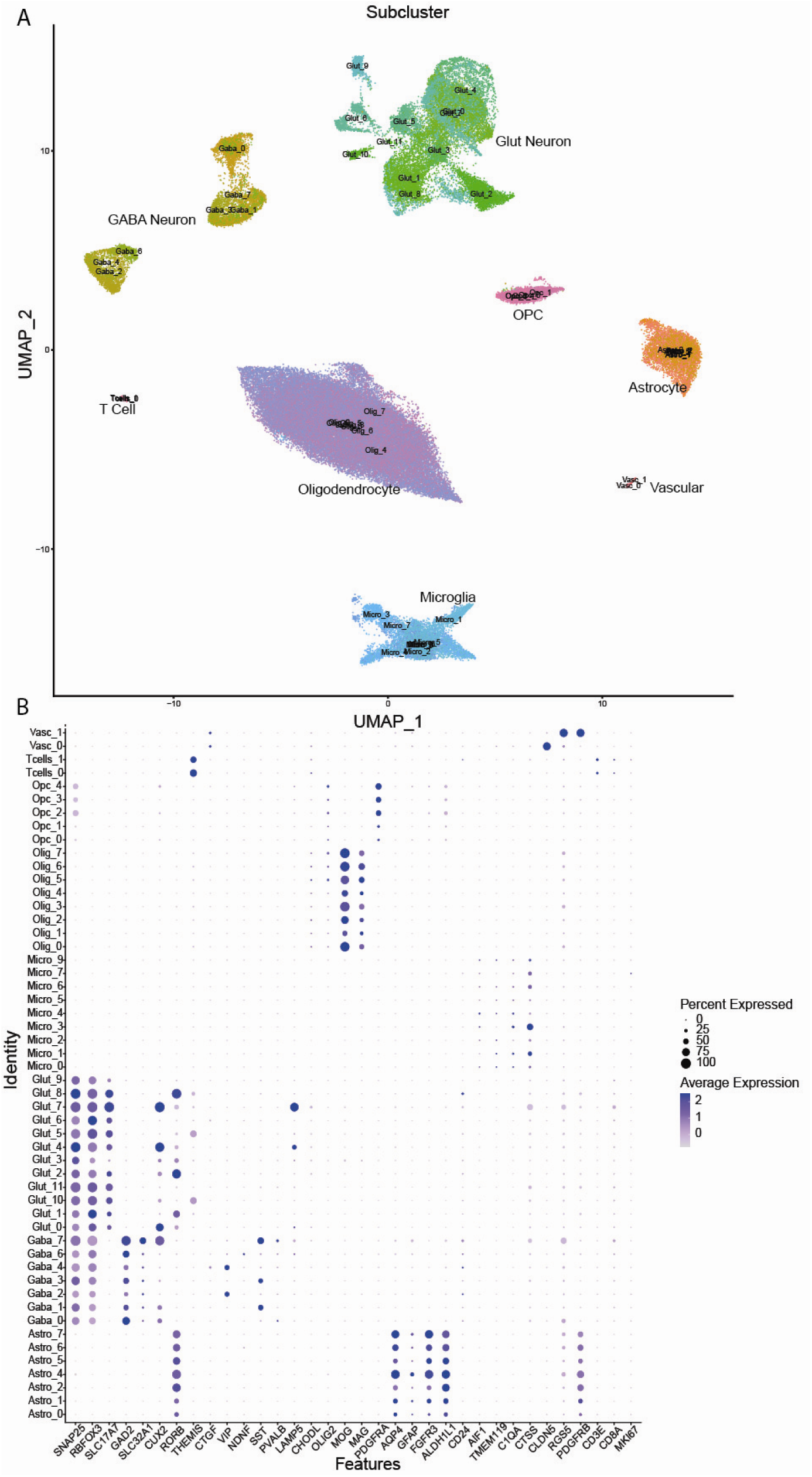
snRNAseq Cell Clustering. **(A)** Uniform Manifold Approximation and Projection (UMAP) shows single-nucleus RNA sequencing of all patient samples (n = 11 patients). Sequencing produced distinct clusters for all major cell types that are clearly separated by single-cell gene expression patterns. **(B)** Gene expression patterns by subcluster, identified by subclustering of broad UMAP clusters. Cell-types for neuronal populations were identified using known gene expression patterns.

We first identified cell proportion counts for each cell-type in the seizure focus and penumbra. In samples from our cohort, we did not find a difference in total neuronal cell count as a fraction of total cells per sample for either excitatory (focus mean: 0.195 ± 0.057, penumbra mean: 0.239 ± 0.027, p = 0.556) (**Supplementary Fig. 2A**) or inhibitory neurons (focus mean: 0.095 ± 0.030, penumbra mean: 0.083 ± 0.022, p = 0.698) (**Supplementary Fig. 2B**). However, when sub-clustering to interneuron cell-types and normalizing to total interneurons per sample, we identified a decreased relative abundance of Parvalbumin (PV) interneurons (PV+ (cluster GABA_0) mean proportion in focus: 0.210 ± 0.047, mean proportion in penumbra: 0.324 ± 0.027, p = 0.049) (**Fig. 4A**) and increased relative abundance of Vasoactive Intestinal Peptide (VIP) interneurons (VIP+ (cluster GABA_2) mean proportion in focus: 0.275 ± 0.041, mean proportion in penumbra: 0.176 ± 0.024, p = 0.099) (**Fig. 4B**) in the seizure focus when compared to the surrounding penumbra. We also see a relative depletion of two putative layer IV/V glutamatergic neuron clusters with expression of RAR-related Orphan Receptor (RORB), as a fraction of total glutamatergic neurons (RORB+ (cluster GLUT_2) mean proportion in focus: 0.060 ± 0.022, mean proportion in penumbra: 0.119 ± 0.021, p = 0.0578) (**Fig. 4C**) (RORB+ (cluster GLUT_8) focus mean: 0.019 ± 0.007, penumbra mean: 0.038 ± 0.009, p = 0.0702) (**Fig. 4D**) in the seizure focus. We did not identify a significant difference in microglia (mean proportion in focus: 0.115 ± 0.049, mean proportion in penumbra: 0.089 ± 0.035, p = 0.192) (**Fig. 4E**), astrocyte (mean proportion in focus: 0.050 ± 0.017, mean proportion in penumbra: 0.072 ± 0.022, p = 0.426) (**Supplementary Fig. 2D**), or oligodendrocyte (mean proportion in focus: 0.480 ± 0.102, mean proportion in penumbra: 0.460 ± 0.105, p = 0.852) (**Supplementary Fig. 2E**) relative cellular abundance in our sequencing data.

**Figure 4.**
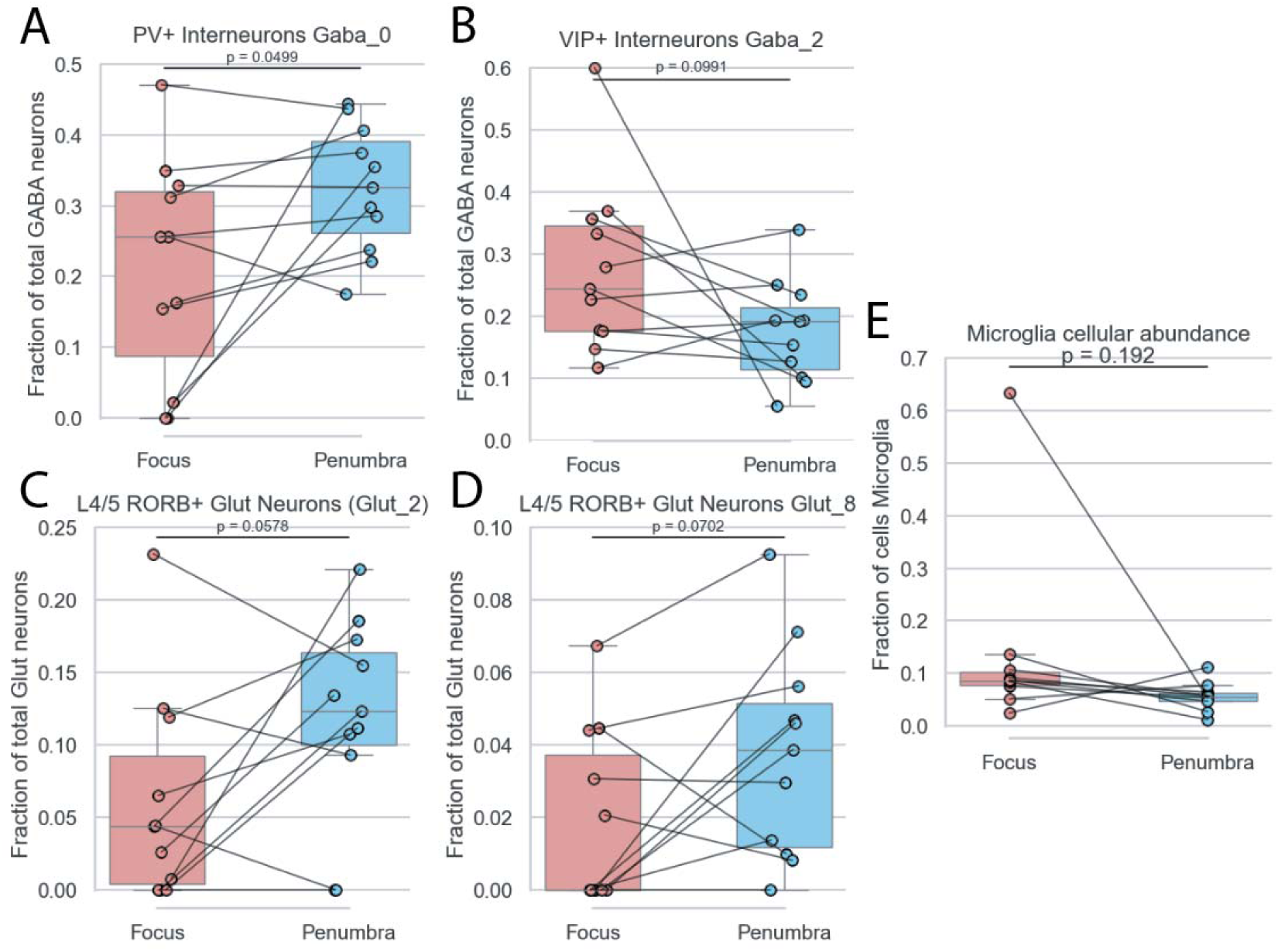
Cellular Abundance via snRNAseq. Cellular abundance of specific cell subcluster populations were calculated as a fraction of the total broad cellular classification (n = 11 patients). **(A)** GABAergic interneurons identified as PV+ interneurons are significantly enriched in the penumbra relative to the seizure focus (p = 0.0499). **(B)** GABAergic interneurons identified as VIP+ interneurons are significantly enriched in the seizure focus, relative to the penumbra (p = 0.0991). **(C)** One subcluster of excitatory neurons identified as RORB+ layer 4/5 glutamatergic neurons are significantly enriched in the penumbra relative to the seizure focus (p = 0.0578). **(D)** A second subcluster of excitatory neurons identified as RORB+ layer 4/5 glutamatergic neurons are significantly enriched in the penumbra relative to the seizure focus (p = 0.0702). **(E)** Clustered microglial cells identified by their broad class identification trend towards enrichment in the seizure focus, but does not reach significance (p = 0.192).

### Immunohistochemical Validation of snRNAseq Findings

We then used IHC to validate key findings seen via snRNAseq, which suggested differential relative abundance of specific neuronal cell types and microglia between the two seizure territories. We identified an increased labeling index (LI) of Iba1+ microglia in biopsies sampled from the seizure focus, relative to the surrounding penumbra (focus mean: 0.196 ± 0.021, penumbra mean: 0.134 ± 0.016, p = 0.007) (**Fig. 5A**).

**Figure 5.**
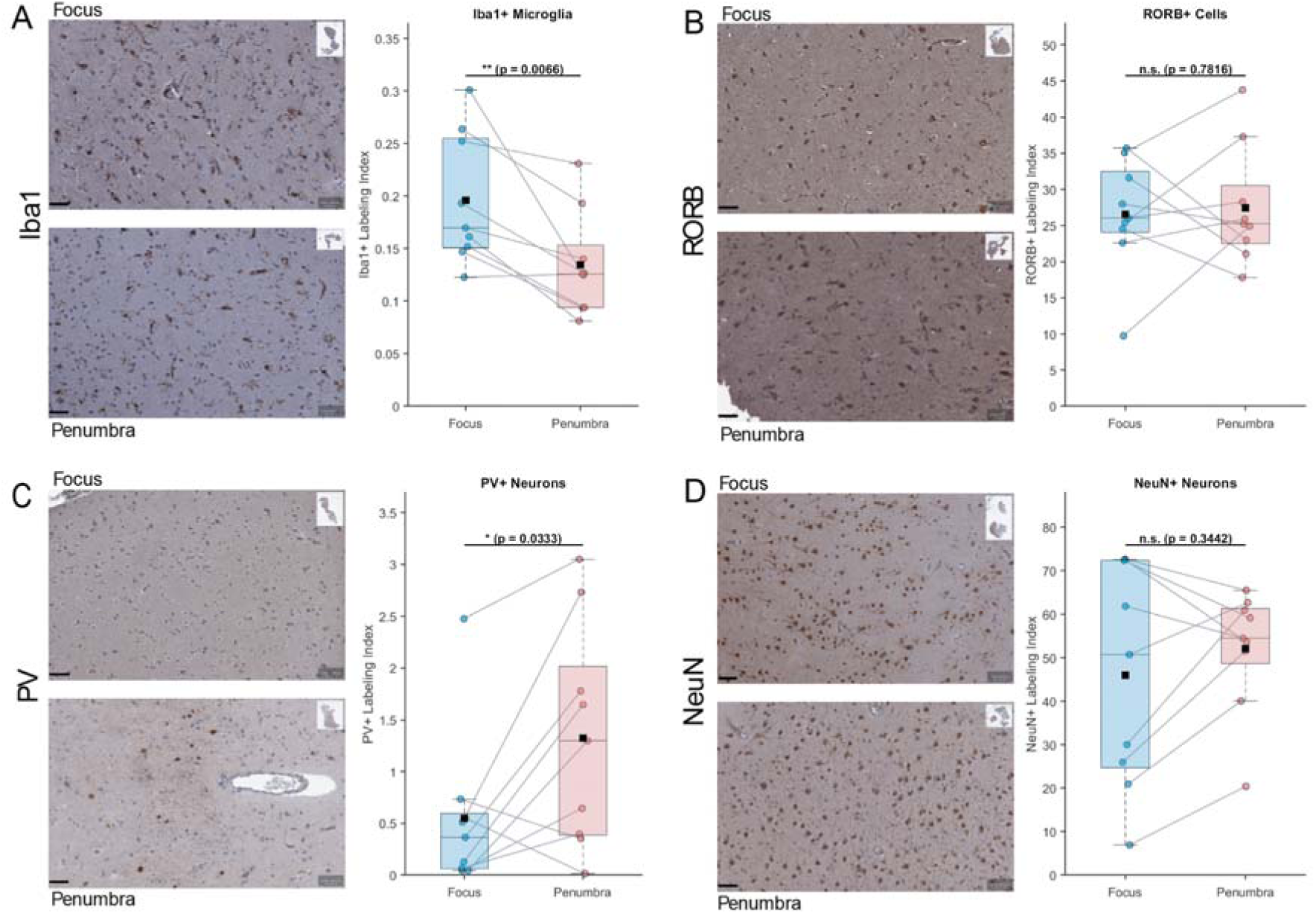
Immunohistochemical Validation of snRNAseq Findings. Cellular labeling indices are calculated as DAB reagent positive cells as a fraction of total hematoxylin-labelled nuclei. **(Left)** exemplary paired tissue IHC micrographs (scale bar 50 µm). **(Right)** Plotted average labeling indices paired per patient (n = 9 patients). We identify **(A)** an increased abundance of Iba1+ microglia in the seizure focus relative to ictal penumbra (p = 0.0066) and **(B)** no relative change in RORB+ cells (p = 0.7816). Of note, we also identify **(C)** a depletion of PV+ interneurons in the seizure focus (p = 0.0333), but **(D)** no corresponding difference in abundance of NeuN+ neurons (p = 0.3442).

When quantifying neuronal labeling indices, we found a relative enrichment in PV+ interneuron labeling index in the ictal penumbra (focus mean: 0.006 ± 0.003, penumbra mean: 0.013 ± 0.004, p = 0.033) (**Fig. 5C**). We found no corresponding difference in total NeuN+ cell labeling index between focus and penumbra samples (focus mean: 0.460 ± 0.085, penumbra mean: 0.521 ± 0.047, p = 0.344) (**Fig. 5D**). This pattern highlights a correlation between PV+ interneurons and the electrographic tissue landscape without corresponding change in overall neuronal abundance.

Validation of RORB+ glutamatergic neurons was performed similarly in cortical regions of tissue collected for histology. However, labeling of RORB+ excitatory neurons yielded no change in abundance of RORB+ cell labeling index (focus mean: 0.265 ± 0.026, penumbra mean: 0.275 ± 0.027, p = 0.782) (**Fig. 5B**). However, we were unable to determine if RORB captured in immunoperoxidase staining was limited to neuronal populations, as this marker has been shown to also be robustly expressed by reactive astrocytes, which was observed in our sequencing data (**Fig. 3B**).^60^

### Consensus Single-Cell Hierarchical Poisson Factorization Analysis

To interpret single nucleus sequencing findings, we used single-cell Hierarchical Poisson Factorization (scHPF) to identify biologically relevant co-expression patterns that demonstrated conserved patterns across patients between the focus and penumbra samples. scHPF is a Bayesian factorization method that identifies both continuous and discrete co-expression patterns from scRNA-seq data in the form of latent factors.^61,62^

The consensus scHPF framework consists of four sequential steps^63^. First, the training data is downsampled to mitigate imbalances in sequencing depth, and genes detected in fewer than 1% of cells are removed. Second, multiple scHPF models are trained across a range of values of *k*, corresponding to different numbers of latent factors. Third, factors derived from these models are clustered to identify stable and reproducible factors across model runs. Finally, the median representations of the stable factor clusters are used as initialization parameters for training the final consensus scHPF model. This contains the recurrent factors as the final value of *k*. Each gene has a score for each factor, quantifying the gene’s contribution to the associated expression pattern. Additionally, each cell assigns a score to each factor, which reflects the contribution of the factor to the observed expression in the cell. scHPF was performed on cell types of interest, including excitatory neurons, inhibitory neurons, and microglia.

By performing paired per-patient analyses of cell scores for each factor, we identified 4 factors that were enriched in excitatory glutamatergic neurons in the ictal penumbra (**Fig. 6**). Factors 0, 5, and 14 were associated with synaptic organization and synaptic signaling gene ontologies. Factor 8 was enriched in ontologies associated with cellular motility and other metabolic alterations. Enrichment of these factors suggests a differential enrichment of synapse-associated gene programs in the penumbra relative to the seizure focus. Other factors identified in excitatory neuronal populations were not differentially expressed in paired analysis (**Supplementary Fig. 3**).

**Figure 6.**
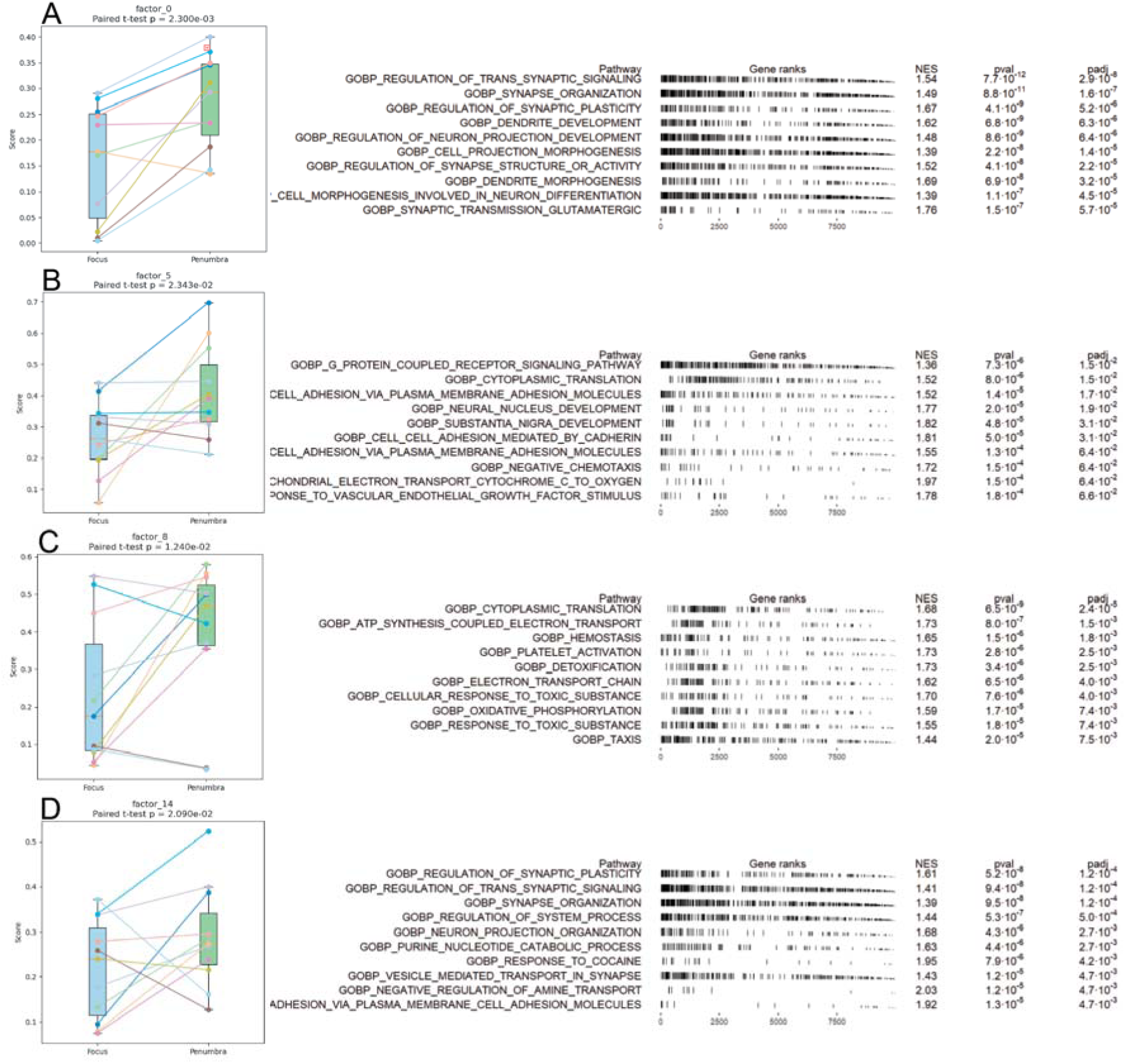
Factorization Model of Excitatory Neuronal Signatures. **(Left)** Average cell score of broad class excitatory neurons per sequenced sample, with lines connecting each patient’s focus and penumbra samples (n = 11 patients), plotted for each individual factor that was identified as significant (p < 0.05 via paired t-test). **(Right)** For each factor that was differentially expressed, an unsupervised GSEA was run using human gene ontologies of biological processes. Top 10 gene ontologies ranked by significance are shown.

Similar analysis of GABAergic inhibitory neurons showed enrichment of 4 factors in the penumbra (Factors 1, 4, 5, and 11), and enrichment of one factor (Factor 13) in the focus (**Fig. 7**). GSEA of biological process ontologies indicated that Factors 1, 5, and 11 were associated with plasticity and neuronal signaling pathways, including dendrite formation and spine development. Factor 4 was associated with various ontologies associated with metabolic and biosynthetic processes. Of note, Factor 13 was differentially enriched in the focus and included leading edge genes associated with synapse assembly. These findings overall suggest a differential enrichment of plasticity markers in inhibitory neurons in the penumbra relative to the seizure focus, consistent with findings in our excitatory neuron populations. Other factors identified in inhibitory neurons were not differentially expressed in paired analysis (**Supplementary Fig. 4**).

**Figure 7.**
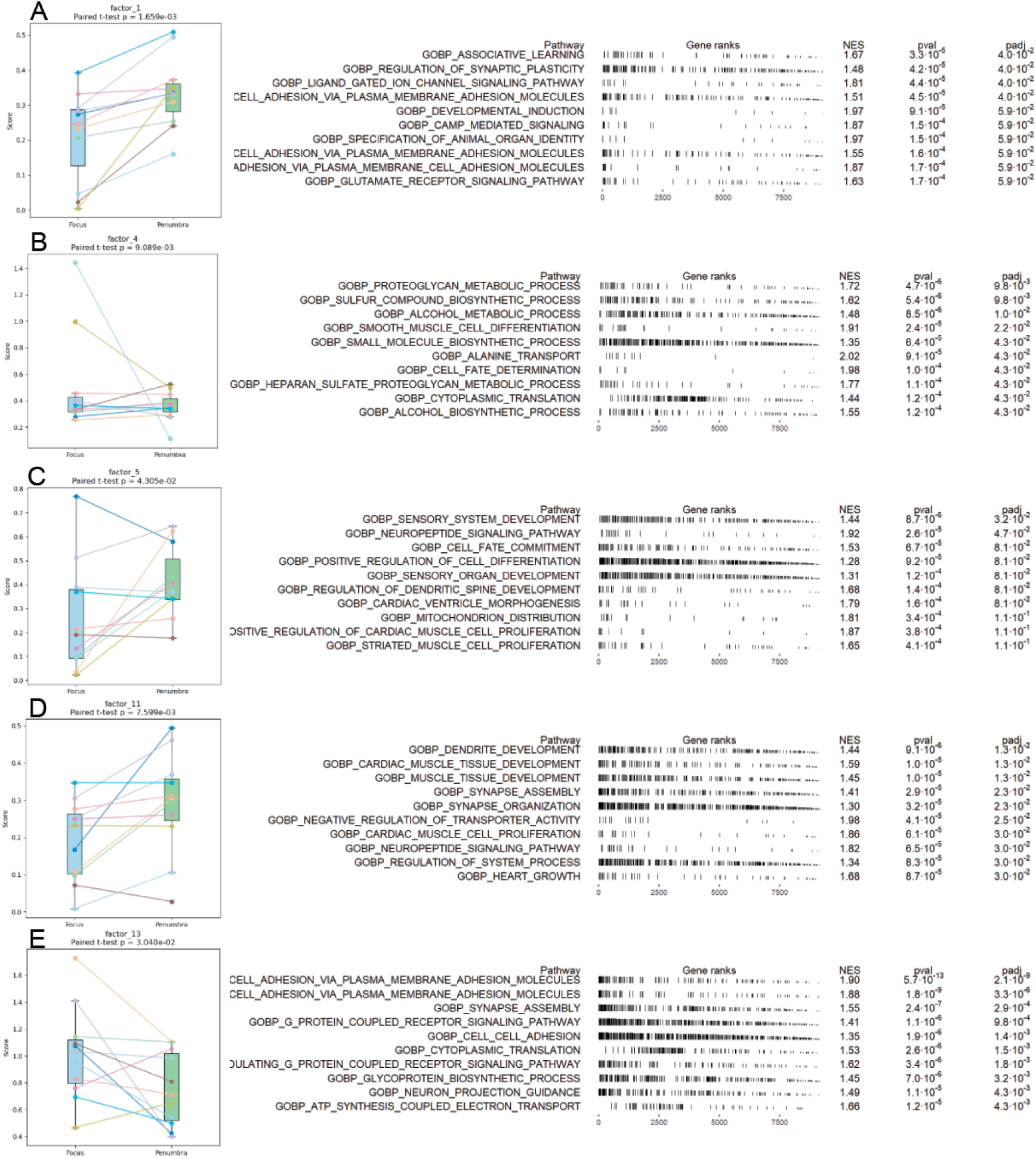
Factorization Model of Interneuron Signatures. **(Left)** Average cell score of broad class inhibitory neurons per sequenced sample, with lines connecting each patient’s focus and penumbra samples (n = 11 patients), plotted for each individual factor that was identified as significant (p < 0.05 via paired t-test). **(Right)** For each factor that was differentially expressed, an unsupervised GSEA was run using human gene ontologies of biological processes. Top 10 gene ontologies ranked by significance are shown.

We observed two microglial factors, one enriched in the penumbra and one in the focus, after removing one patient from the factorization model due to it not passing quality control metrics (**Fig. 8**). However, gene ontologies of biological processes did not identify consistent gene programs with interpretable, biologically relevant signals. Other factors identified in microglia were not differentially expressed in paired analysis (**Supplementary Fig. 5**).

**Figure 8.**
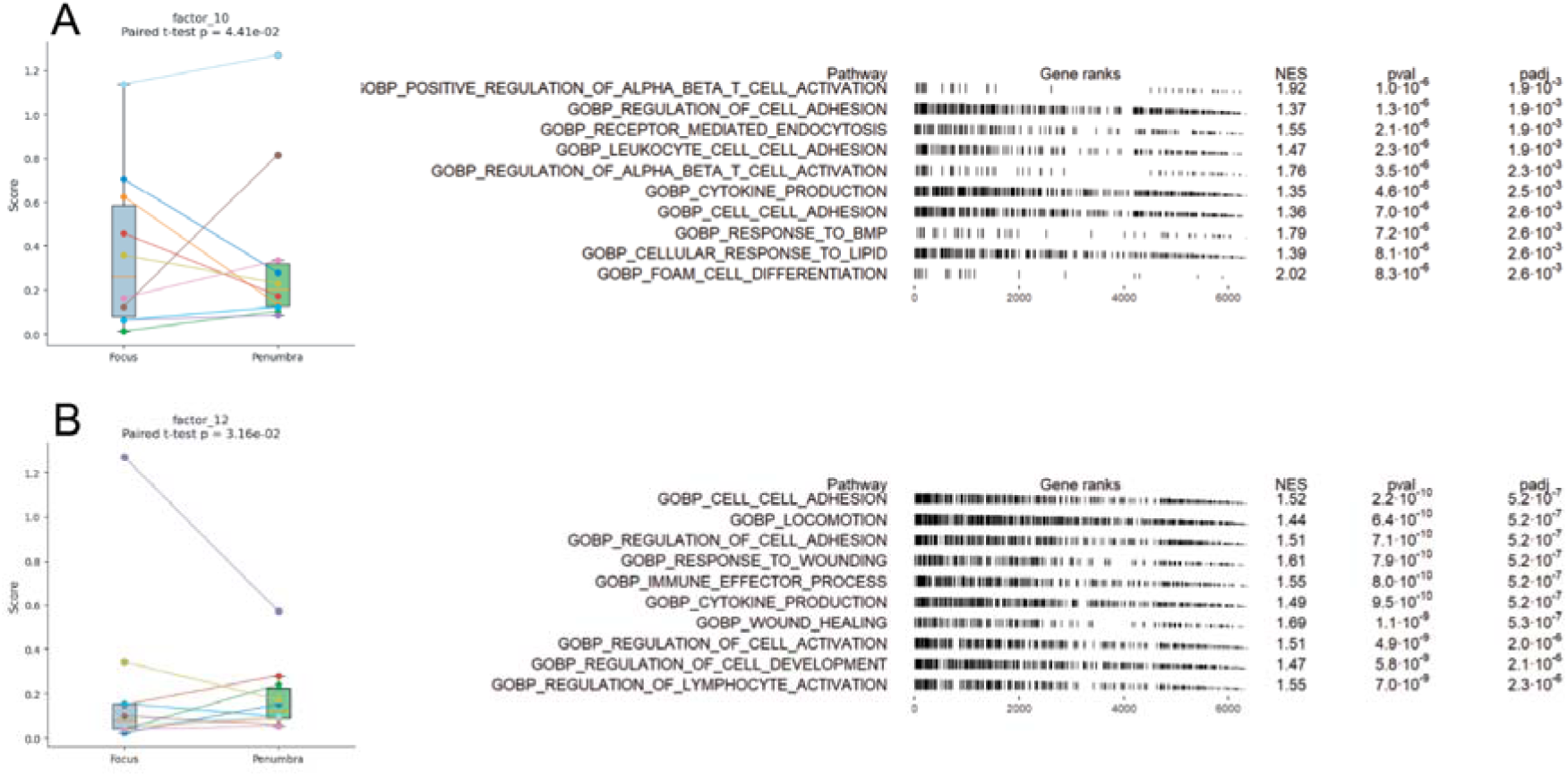
Factorization Model of Microglial Signatures. **(Left)** Average cell score of broad class microglia per sequenced sample, with lines connecting each patient’s focus and penumbra samples (n = 11 patients), plotted for each individual factor that was identified as significant (p < 0.05 via paired t-test). **(Right)** For each factor that was differentially expressed, an unsupervised GSEA was run using human gene ontologies of biological processes. Top 10 gene ontologies ranked by significance are shown.

## Discussion

### Overview

In this study, we use a novel methodology to sample distinct electrophysiologic territories of seizure involvement in patients with intractable focal epilepsy and identify distinct cellular and molecular transcriptional patterns across the electrophysiologic seizure landscape. We identify a landscape of pathologic tissue features that co-occur with electrophysiologic features of the tissue during seizure activity. Our sampling method provides a unique view into conserved tissue-based readouts of these regions, and we discovered conserved findings that emerged despite the mix of epilepsy pathologies, patient age, and duration of epilepsy which have been historically used to stratify sequencing and histological analyses.^64^

Tissue-based analyses in focal epilepsies have historically been limited by lack of a comparator, as sampling otherwise healthy tissue is limited to post-mortem autopsies or distinct brain pathologies, confounding comparisons. We therefore compare paired biopsies sampled from the same individual, both invaded by seizure activity, but that are distinguished by EEG features reflecting their involvement in seizure activity. We differentiate regions involved in seizure onset as opposed to territories being subjected to synaptic current distribution but restrained localized neuronal firing, which exhibit a low likelihood of active seizure invasion. Our methodology thus provides a substantive advance in identifying molecular and cellular features that underlie these distinct electrographically defined regions.

We identify changes in neuronal abundance with corresponding tissue immunohistochemistry, differential gene expression patterns, and show differential enrichment of gene expression programs identified through scHPF. This advanced sequencing analysis produced a study-specific set of transcriptional patterns that allows us to analyze changes in transcriptional programs of biological relevance to our dataset. Our findings highlight possible tissue features that predispose ictal involvement and seizure generation or features that represent downstream tissue responses and compensatory mechanisms to pathological ictal activity. Given the inclusion of various neocortical epilepsy etiologies across a broad range of patient age, seizure duration, and involved brain region, we additionally inherently capture variability in cell abundance and transcriptional state. However, our identification of consistent signatures that exhibit significance when using paired analyses by patient reflect broader consistent differences in tissue markers found in the focus and penumbra across focal epilepsies and validate our sampling methodology.

### Plasticity Associated with Focal Seizures

Our analyses identified multiple co-expressed factors in both excitatory and inhibitory neuronal populations that show a transcriptional upregulation of genes associated with synaptic remodeling differentially enriched in the ictal penumbra. Neuroplasticity allows for constant remodeling of neuronal microcircuitry, allowing for changes in structure, density, and strength in signal transmission in specific neuronal connections.^65^ These changes have been hypothesized as both direct remodeling due to proximal neuronal activity and as potential compensatory mechanisms that conserve excitatory-inhibitory interactions.^66^ This structural remodeling due to proximal pathologic activity has also been hypothesized as a mechanism for expansion of epileptogenic networks into otherwise healthy circuitry.^67^ Conversely, structural remodeling has similarly been proposed as a predictor of clinical response to responsive neurostimulation (RNS), due to observed correlation of functional connectivity with seizure reduction in long term follow-up.^68^ Taken in this context, our findings show the ictal penumbra as a region of relative elevated plasticity-associated signatures, characterizing this region as a reactive and dynamic landscape in comparison to the seizure focus and a potent substrate for site-specific remodeling for long-term seizure control.

We are limited in our ability to make a distinction as to whether this plasticity is a driver of long-standing network recruitment or whether these are reactive changes to combat repetitive excitatory input or some combination of these phenomena. While we interpret the differential changes in synaptic plasticity to represent an enrichment in the ictal penumbra, our results could also plausibly be interpreted as a depletion of synaptic transcriptional signatures in the focus due to the nature of our sample comparison. However, our results do support focal epilepsies as a dynamic set of diseases that exhibit ongoing pathologic changes to healthy cortex that over time may break down excitatory-inhibitory interactions in the ictal penumbra and promote recruitment to local ictogenic networks.

There is ongoing debate on the long-term consequences of epilepsy disorders and seizure activity on the functional activity of the brain.^69–71^ Following focal epileptiform activity, intrinsic mechanisms of neuronal plasticity aimed at maintaining homeostasis can remodel brain circuitry to reinforce and potentially promote tissue hyperexcitability.^72^ Proinflammatory cytokines, reactive astrocytes, and reactive microglia are consistent features of epilepsy-induced inflammation that have been identified from resected tissue from patients undergoing epilepsy surgery.^73,74^ This seizure-induced inflammation has been suggested as yet another pro-epileptogenic mechanism, with inflammatory cytokines and elevated complement having been shown to exacerbate seizure generation.^75,76^ Further study of neuronal circuitry and the downstream reactive immune compartment in epileptic regions at a tissue-based level may aid in highlighting epilepsy-related inflammation as a target for therapeutic intervention.^77^

### Immune drivers and reactions in epilepsy

In our cohort, we identified a significant enrichment in microglial labeling index in the seizure focus via IHC. Immune localization has been identified consistently in regions of seizure involvement in both preclinical mouse models and patient samples. Indeed, microglia are now known to play a critical role in shaping neuronal architecture and modulate neuronal networks in developmental states, potentially playing a pivotal role in early disease pathology.^8,78^ Excitatory neuron bursting and excitotoxicity has been shown to release immune chemoattractants including purinergic signals and other damage-associated molecular patterns, or DAMPS.^78^ Prior literature has further suggested inflammatory processes contribute to local ictogenesis through a number of methods including remodeling of neuronal architecture, connectivity, and alterations to the extracellular environment.^79^

We earlier discuss an alternative interpretation of our results implicating a relative depletion of synaptic signatures in the seizure focus. Such a finding, paired with our finding of increased microglial localization to the seizure focus via IHC, may relate closely to prior literature that highlights microglia as central players in reshaping neuronal synapses, mediated by complement-driven synaptic pruning. Aberrant synaptic engulfment by reactive microglia has recently been implicated in synaptic dysfunction seen in epilepsy disorders.^79–82^ This process appears to be tightly regulated in healthy cortex, with both excessive and deficient synaptic pruning predisposing animal models to developing seizures.^80,83,84^ Recent studies further suggest that microglia may preferentially engulf inhibitory synapses during epileptogenesis, disrupting the excitatory/inhibitory balance and promoting neuronal hyperexcitability.^79,83^ Further work should determine a causal relationship between microgliosis and synaptic pruning in disrupting excitatory-inhibitory network dynamics.

### Neuronal cell-type involvement and susceptibility in epilepsy

Interneurons critically allow for network activity to perform complex computational activity and dynamic stimulus responses without leading to recurrent network excitation.^85^ Parvalbumin-expressing basket interneurons in particular mediate network inhibition in cortical regions, therefore playing a critical role in setting the excitatory-inhibitory balance of local networks and maintaining physiologic network excitability.^85,86^ In our samples, we identified a significant differential decrease in PV+ interneurons in the seizure focus via both snRNAseq and IHC. Pathology in PV+ interneurons is consistent with existing computational and experimental animal models of neocortical epilepsy.^17^ The loss of PV interneurons and their function has been previously noted in hippocampal and neocortical samples from temporal lobe epilepsy and epilepsy associated with cortical malformations, although results were limited by lack of electrophysiology-guided sampling and single-cell resolution.^43–45^ However, changes in cellular abundance may not inherently provide conclusive evidence of functional consequences towards ictogenesis.

This observed difference in PV+ interneuron cellular abundance supports a hypothesis that regions deficient in PV+ interneurons are predisposed to high-intensity epileptiform activity and impaired inhibitory restraint, permitting seizure initiation and propagation when the restraint fails.^12,17^ This cellular compositional change could alternatively point towards a susceptibility of these cells towards cell death in response to local high-amplitude synchronous currents simply due to their small size, which could provide insight into hyperexcitability and cognitive dysfunction across a wide range of disease.^87^ Further study should assess the effects of local PV+ interneuron loss in neocortical networks to understand functional consequences on regional network activity.

### Deep Layer Excitatory Neuron Involvement in Agranular Cortex

snRNAseq cell abundance analysis revealed a significant differential enrichment of two clusters identified as RORB expressing pyramidal neurons in penumbral samples. RORB+ excitatory neurons can be found largely in the granular cortex, or layer 4/5 of mammalian neocortical tissue. Prior work has demonstrated significant involvement of deep cortical layers, including the granular cortex and infra-granular cortex, in seizure generation and onset.^88^ These initiating events are often followed by current flow and synaptic spread mediated through supra-granular layers. Studies have suggested that the granule layer is a major player in seizure propagation and driving seizure spread in neocortical epilepsy in somatosensory cortex. However, the absence of granular cortex in large swaths of the frontal lobe including motor cortex suggests alternative mechanisms for onset and spread of neocortical seizures.^89^ Our identification of decreased RORB+ neuronal abundance in the seizure focus supports literature pointing to involvement of these deep layer neurons as key players in ictogenesis and seizure pathology.

### Conclusions and Clinical Implications

Our study provides a novel methodology that allows for direct comparison between the seizure focus and ictal penumbra. Our findings provide the first in-patient direct comparisons of transcriptional and tissue-based readouts that provide insight into a heterogeneous electrophysiologic landscape. We identify neuronal compositional and transcriptional signatures that differentiate the seizure focus and penumbra that directly correlate with their involvement in seizure activity. Further work will need to translate localized tissue and biophysical functional differences to their effects on large-scale network activity which may appear coordinated, but which may rely on this heterogeneous landscape of functional connectivity across seizure territories.

While conventional therapeutic interventions for intractable epilepsy have relied on tissue resection, non-destructive neuromodulation approaches could apply site-specific neuromodulation to the heterogeneous seizure landscape for prolonged seizure control.^90^ If our plasticity findings are functionally validated, this may suggest the ictal penumbra as a target for structural reorganization via site-specific RNS for long-term seizure control.^91^ Changes in interneuron abundance could relate to experimental therapies which have shown implantation of iPSC-derived interneurons can increase local inhibitory tone in ictogenic networks with promising preclinical efficacy, now in early clinical testing (NCT05135091).^92–94^ Ultimately, our findings demonstrating an enrichment of synaptic plasticity ontologies in the penumbra, paired with our findings of immune infiltration in ictogenic regions, support a broader hypothesis of focal epilepsies as dynamic pathologies that exhibit a heterogeneous landscape of tissue features that correlate with electrographic involvement.

## Acknowledgements

We would like to thank the clinical epileptologists and fellows at Columbia University Irving Medical Center. We would also like to thank the neurosurgeons and residents who were involved in patient care. We would especially like to thank the patients involved in the study, without whom this work would not be possible. Their selfless contributions allow for advances in future clinical care and research advances. We thank the staff at the Molecular Pathology Shared Research Core at Columbia University Irving Comprehensive Cancer Center for help embedding and sectioning tissue. This work was also supported by the Columbia University Medical Center Cancer Center.

## Supplementary material

Supplementary material is available at *Brain* online.

