## Supplementary Figures for "Electrophysiologically Targeted Biopsies Reveal the Transcriptional Landscape of Focal Epilepsy"

**Paired Biopsies of Seizure Territories in Focal Epilepsy Reveal Distinct Cellular Signatures**

710 West 168th Street,

New York,

NY 10032 USA

**Running title**: Paired Biopsies of Seizure Territories

**Keywords:** focal epilepsy, seizure focus, ictal penumbra, paired biopsy, seizure localization

**Supplementary Figures**

**Supplementary Figure 1:** UMAP clustering by patient sample and seizure territory

**Supplementary Figure 2:** Cellular abundance of broad class cell clustering

**Supplementary Figure 3:** Excitatory Neurons factor model, all factors

**Supplementary Figure 4:** Inhibitory Neurons factor model, all factors

**Supplementary Figure 5:** Microglia factor model, all factors

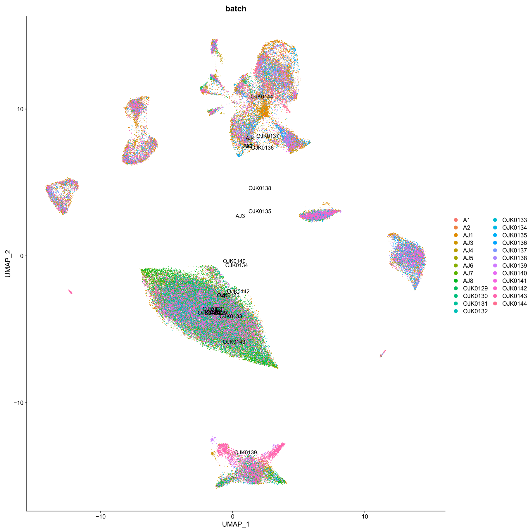

A

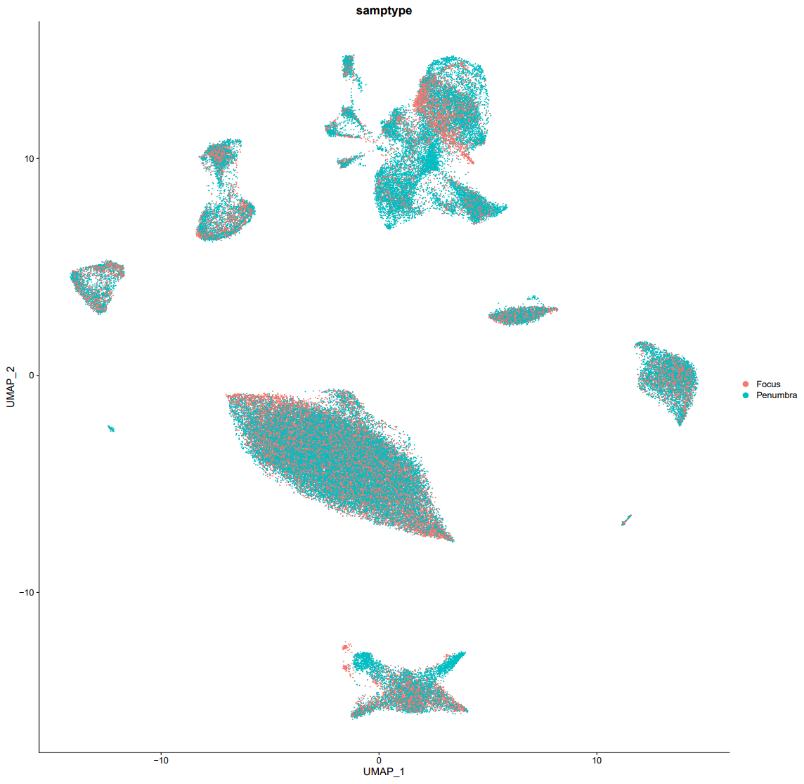

B

**Supplementary Figure 1: UMAP clustering by patient sample and seizure territory.** *These plots help visualize the distribution of cells in clustering analysis and demonstrate the validity of our sequencing quality control.* ***(A)*** *UMAP depicting sample distribution, each color representing a unique biopsy sample. All samples contained cells in all broad cell classes.* ***(B)*** *UMAP demonstrating distribution of cells from Focus (Orange) and Penumbra (Blue), demonstrating all subclusters are well represented by both seizure territories.*

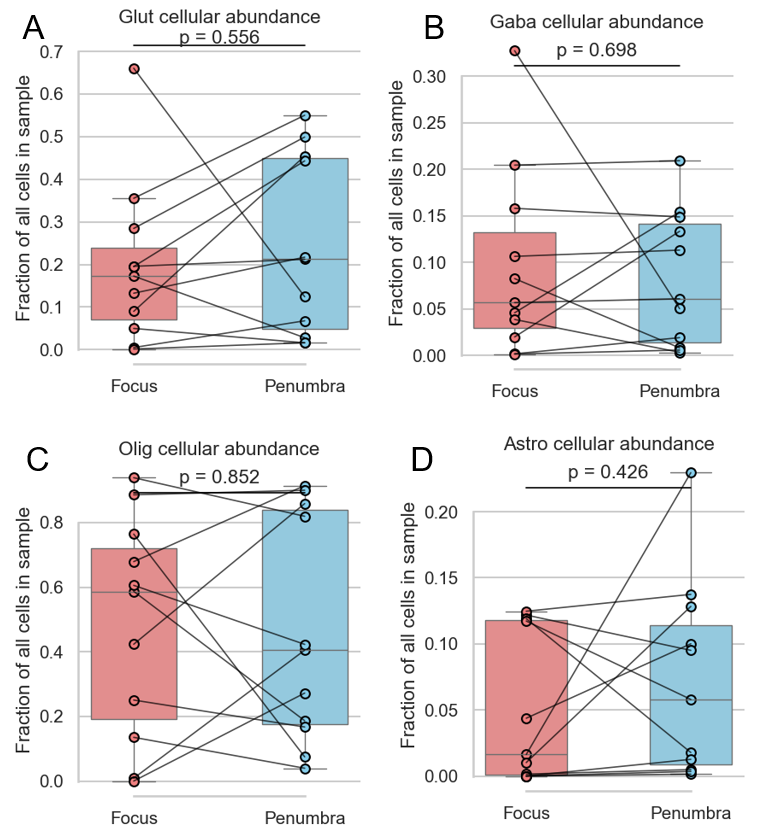

**Supplementary Figure 2: Cellular abundance of broad class cell clustering.** *Paired comparisons of broad cell classification demonstrates no difference in broad cell class abundance. Abundance was normalized to total cell count from all broad classes within each sample. This is shown for* ***(A)*** *Excitatory neurons* ***(B)*** *Inhibitory neurons* ***(C)*** *Oligodendrocytes and* ***(D)*** *Astrocytes.*

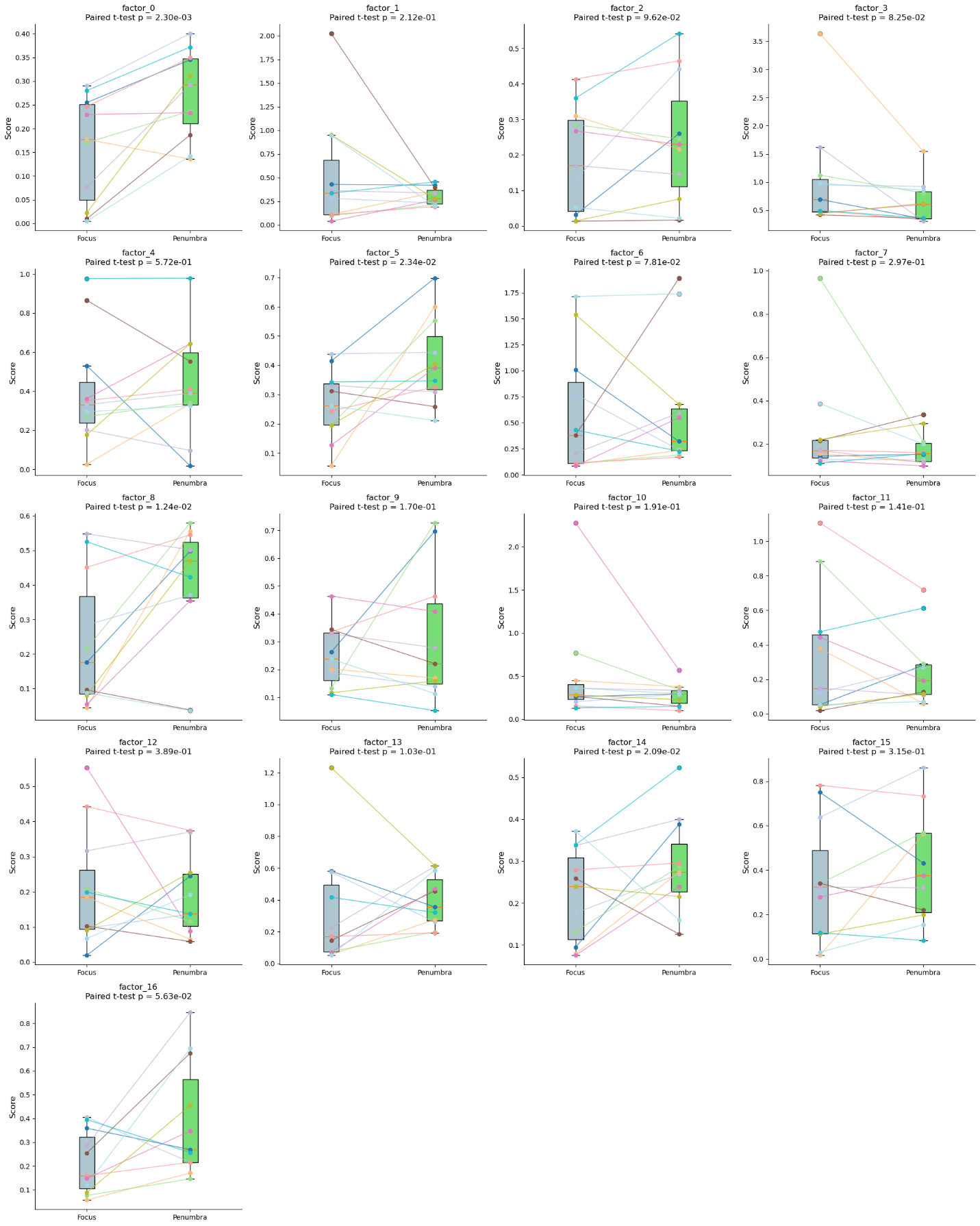

**Supplementary Figure 3: Excitatory Neurons factor model, all factors.** *Each factor derived from the excitatory neuronal scHPF model resulted in an average cell-score per patient sample. Focus samples were compared to penumbra samples in a pairwise analysis.*

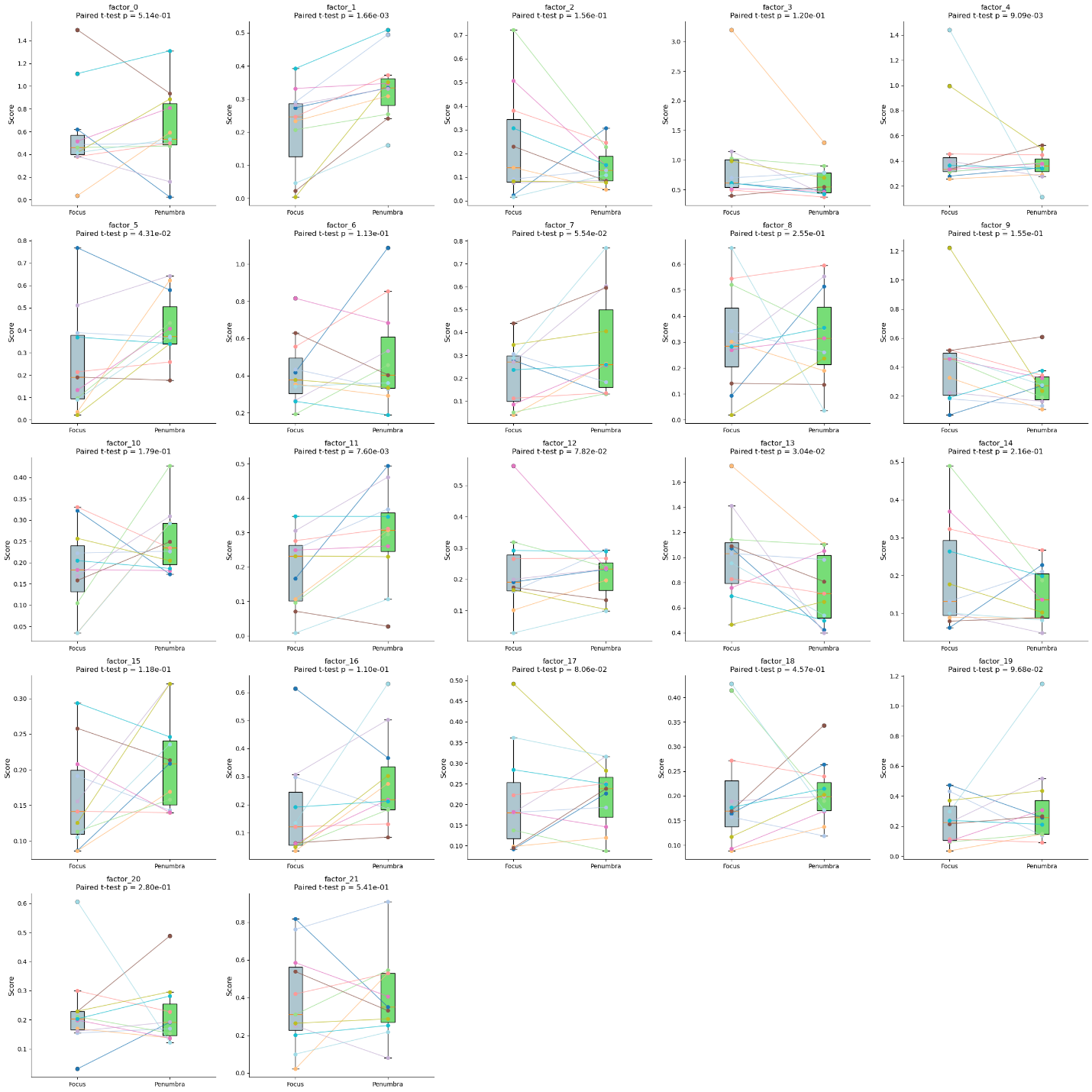

**Supplementary Figure 4: Inhibitory Neurons factor model, all factors.** *Each factor derived from the inhibitory neuronal scHPF model resulted in an average cell-score per patient sample. Focus samples were compared to penumbra samples in a pairwise analysis.*

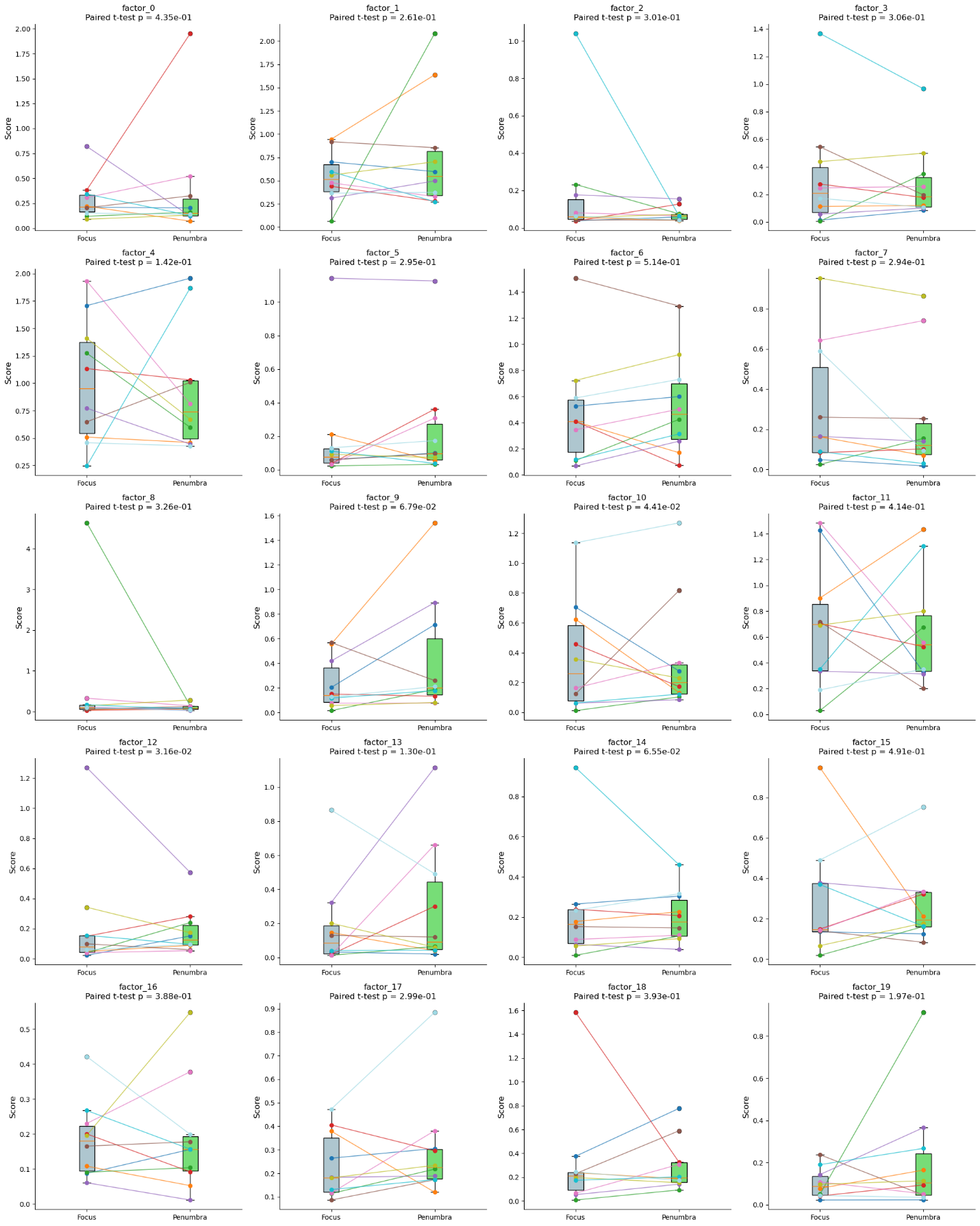

**Supplementary Figure 5: Microglia factor model, all factors.** *Each factor derived from the Microglial scHPF model resulted in an average cell-score per patient sample. Focus samples were compared to penumbra samples in a pairwise analysis.*
